# Suppression of the dihydrolipoamide dehydrogenase gene (*dld-1*) protects against the toxicity of human amyloid beta in *C. elegans* model of Alzheimer’s disease

**DOI:** 10.1101/228429

**Authors:** Waqar Ahmad

## Abstract

Declines in energy metabolism and associated mitochondrial enzymes are linked to the progression of Alzheimer’s disease (AD). Dihydrolipoamide dehydrogenase (*dld*) and two of its enzyme complexes namely, pyruvate dehydrogenase and α-ketoglutarate dehydrogenase are associated with AD and have a significant role in energy metabolism. Interestingly, *dld* gene variants are genetically linked to late-onset AD; and reduced activity of DLD-containing enzyme complexes has been observed in AD patients. To understand how energy metabolism influences AD progression, we suppressed the *dld-1* gene in *C. elegans* expressing the human Aβ peptide. *dld-1* gene suppression improved many aspects of vitality and function directly affected by Aβ pathology in *C. elegans.* This includes protection against paralysis, improved fecundity and improved egg hatching rates. Suppression of the *dld-1* gene restores normal sensitivity to aldicarb, levamisole and serotonin, and improves chemotaxis. Suppression of *dld-1* does not decrease levels of the Aβ peptide, but does reduce the formation of toxic Aβ oligomers. The mitochondrial uncoupler, carbonyl cyanide 4-(trifluoromethoxy)phenylhydrazone (FCCP) acts synergistically with Aβ to overcome the protective effect of *dld-1* gene suppression. Another metabolic toxin, phosphine, acted additively with Aβ. Our work supports the hypothesis that lowering energy metabolism may protect against Aβ pathogenicity, but that this may increase susceptibility to other metabolic disturbances.

## 1. Introduction

One of the main pathological hallmarks of AD that underlies the neuronal dysfunction and dementia is extracellular accumulation of amyloid beta (Aβ) plaques resulting from protein misfolding [1]. In addition to the accumulation of Aβ, neuroimaging studies of AD brains found induced aerobic glycolysis in areas severely affected by the disease at the preclinical stage, but reduced glucose metabolism and diminished activities of mitochondrial enzymes at latter stages of the disease [2-5]. The cause and effect relationships between these observations is unclear as impairment of energy metabolism may induce protein misfolding, leading to formation of Aβ plaques, but the opposite may also be true as production and accumulation of Aβ may also damage energy metabolism [6-12].

The decrease in energy metabolism with age has been interpreted in two opposing ways; as a main cause of AD, or as a protective response against the symptoms of the disease. The first interpretation is mostly supported by studies conducted at a late stage of the disease on postmortem brains, making it difficult to assign causality [13-17]. In contrast, recent studies on mixed stage AD samples support that the down-regulation of energy metabolism is a protective factor, leading to the hypothesis that a decrease in nutrient and oxygen supply minimizes neural activity, thereby decreasing the repair burden [18, 19]. This is supported by results from a transgenic mouse model of AD in which upregulation of aerobic respiration is clearly harmful [20].

The difficulty in understanding the role of metabolic decline on AD relates to the inaccessibility of AD affected brains during progression of the disease. This situation makes it difficult to distinguish cause from consequence and necessitates reliance on AD disease models. The presence of Aβ in mice and neuronal cell lines causes mitochondrial dysfunction by inducing mitochondrial DNA damage, disrupting mitochondrial redox potential and inducing oxidative stress [21-24]. Exposure to 2-deoxy-d-glucose (2DOG), which is an analogue of glucose that inhibits both glycolysis and oxidative phosphorylation, causes a decrease in Aβ toxicity [25, 26]. Both of these results could be interpreted as lower rates of glucose metabolism having a protective effect. Furthermore, induction of oxidative phosphorylation with dichloroacetic acid (an activator of pyruvate dehydrogenase (PDH)) increases neuronal sensitivity toward Aβ [27]. These findings reinforce the interpretation that there is a positive correlation between the rate of energy metabolism and Aβ toxicity.

Increased risk of late-onset AD is genetically linked to the human *dld* locus [28]. The DLD enzyme is a subunit of three ketoacid dehydrogenase complexes, each of which contributes to energy metabolism, pyruvate dehydrogenase PDH, α-ketoglutarate dehydrogenase complex (KGDH) and branched chain ketoacid dehydrogenase complex (BCKDH) [29, 30]. A decrease in the activity of PDH and KGDH is associated with neurodegeneration as reduced activities of these complexes are observed in post-mortem brain tissues and fibroblasts of patients with either Alzheimer’s or Parkinson’s disease [31-36]. Repressed activity of the KGDH complex, in particular by creating mice that are heterozygous for a knockout mutation of one of its subunits (E2), decreases glucose utilization in the cortex, which mimics the situation in cortical neurodegenerative diseases [37, 38]. Targeted disruption of DLD can also reduce the activities of KGDH and PDH in mice [39], which in the case of PDH blocks the connection between glycolysis and the TCA cycle. [40, 41]. Thus, a direct link between DLD activity and AD progression is a distinct possibility.

To explore the relationship between metabolism and AD, we suppressed *dld-1* in the nematode *C. elegans* that express human Aβ. *C. elegans* is well-suited for such studies as it has been used extensively to study the genetics of aging and associated age-related diseases such as AD. Well-developed models exist in which human Aβ is expressed in either muscle or neuronal cells of *C. elegans* [42, 43]. These strains can be used to monitor the influence of metabolic changes on the aggregation of Aβ peptide either directly or through altered behaviour resulting from cellular malfunction. Assays that have been developed include progressive paralysis, a decrease in chemotactic ability and altered response to neurotransmitters. These assays allow convenient visual monitoring of the molecular processes leading up to Aβ aggregation. A decrease in Aβ mediated pathology in response to suppression of *dld-1* supports the notion that decreased energy metabolism is neuroprotective.

## 2. Materials and Methods

### 2.1. Nematode strains

*C. elegans* strains used in this study are the wild type strain, N2 (Bristol), and the long-lived, stress resistant *dld-1* mutant, *dld-1*(*wr4*). *dld-1*(*wr4*) strain contains a A460V missense mutation and showed resistance against phosphine exposure that can also be achieved using *dld-1* RNAi in wild type [44, 45]. Strains expressing human β-amyloid peptide in muscle cells include CL2006 (dvIs2 [pCL12(unc-54/human Abeta42 minigene) + pRF4]), which produces the human Aβ peptide constitutively and CL4176 (dvIs27 [(pAF29)myo-3p::Abeta (42)::let 3'UTR) + (pRF4)rol-6(su1006)]), in which Aβ peptide is produced when the temperature is increased from 16°C to 23°C. The strain CL802 [smg-1(cc546) I;rol-6(su1006) II] was used as a control for CL2006 and CL4176 in assaying paralysis/ movement. The use of these strains as a worm model of AD was documented previously [46]. We used strain CL2355 (dvIs50 [pCL45 (snb-1::Abeta 1-42::3' UTR(long) + mtl-2::GFP] I), in which Aβ is expressed pan neuronally, to complement studies on the strains in which Aβ was expressed in muscle cells. The control strain for CL2355 was CL2122 (dvIs50 [(pD30.38) unc-54 (vector) + (pCL26) mtl-2::GFP] I) [43, 46-48]. Sod-3::GFP reporter strain CF1553 (muIs84 [(pAD76) sod-3p::GFP + rol-6(su1006)) was also used in this study.

### 2.2. Culture conditions

Mixed-stage cultures of *C. elegans* were maintained on nematode growth medium (NGM) seeded with *E. coli* OP50 at 20°C, except strains CL4176 and CL2355, which were maintained at 16°C to suppress Aβ expression. Synchronised cultures for bioassays were obtained by standardized protocols described previously [43, 46-48]. Wild type, *dld-1*(*wr4*) mutant and Aβ transgenic worms CL4176 were all initially cultured at 16°C for 36 hours after which the temperature was increased to 23°C for 36 hours except for the paralysis assay for which the temperature was further increased to 25°C to maximise expression of the Aβ transgene. Phenotypes of the worms were monitored by visual observation under a microscope and/or quantified using the WormScan procedure [49].

### 2.3. dld-1 gene suppression by RNAi

The *E. coli* strain SJJ_LLC1.3 (Source Bioscience), which expresses double-stranded RNA of the *dld-1* gene was fed to each of the four *C. elegans* strains to suppress expression of the *dld-1* gene [50]. Briefly, the bacteria were cultured in LB medium containing 100 μg/mL ampicillin overnight with shaking at 37°C. 300 μL of this bacterial culture was transferred to NGM plates containing 100 μg/mL ampicillin and 1 mM IPTG. The plates were incubated at 25°C overnight to allow the bacteria to grow. Synchronised L1 worms were transferred to the bacterial plates and kept at 16°C for 36 hours. After a further 36 hours at 25°C the worms were ready for use in the assays described below. Mock gene suppression controls were treated in exactly the same way except that the bacterial strain (HT115) for the controls contained the plasmid vector without the *dld-1* gene fragment.

### 2.4. Paralysis and mortality assays

Paralysis can be defined as time-dependent observable decrease in worm’s muscular movement. At severe stage it may lead to complete halt of movement and worms might be consider as dead. Paralysis could be slow down or reversible. [51]. Mortality could be defined as acute death after certain period of time due to decline of cellular functions and organelles. Synchronised, L1 stage worms were transferred to NGM plates that had been seeded with the *dld-1* RNAi *E. coli* or an *E. coli* strain (HT115) containing an empty vector as a control. After 36 hours at 16°C, worms were upshifted to 25°C. The worms were then scored for paralysis every second hour after an initial 24-hour period until the last worm became paralysed. For mortality assays, worms were counted as dead or alive after treatment.

### 2.5. Touch response assay

Touch response assays were performed at 20°C on synchronised L4 worms after inducing Aβ expression at 23°C as described for *dld-1* gene suppression. Fifteen (15) animals of each strain were selected arbitrarily and put on freshly made NGM plate. Worms were left on plates for 2 minutes to allow them to equilibrate to the new conditions. Worms were then touched on the head or tail region using a platinum wire to stimulate locomotion and body bends were then counted for 30 seconds.

### 2.6. Aldicarb and levamisole assays

Worms prepared as described for *dld-1* gene suppression were incubated in the presence of 1 mM aldicarb, an acetylcholinesterase inhibitor [52]. In parallel with the aldicarb experiment, we also exposed worms to 0.2 mM levamisole, a cholinergic agonist [53]. The number of active worms was counted every hour until all worms became paralyzed.

### 2.7. Phosphine exposure assay

Nematodes were fumigated with phosphine at 500 ppm and 2000 ppm as described previously [54]. Briefly, a synchronised population of 48 hour old (L4) nematodes was washed with M9 buffer and approximately 80-100 nematodes were transferred to each well of 12-well tissue culture plates containing 2.5 mL of NGM agar per well pre-seeded with either empty vector strain *HT115* or *E. coli* strain SJJ_LLC1.3. Nematodes were exposed to phosphine for 24 hours in glass fumigation chambers. The chambers were opened and worms were allowed to recover for 48 hrs in fresh air with lids on plates. The numbers of surviving nematodes were then counted.

### 2.8. 5-HT sensitivity assay

To determine the level of Aβ-induced 5-HT hypersensitivity, serotonin (creatinine sulfate salt) was first dissolved in M9 buffer to 1 mM as described previously [43]. Synchronized worms were then washed with M9 buffer and transferred into 200 μl of the 1 mM serotonin solution in 12-well assay plates. The worms were scored as either active or paralysed after 5 minutes.

### 2.9. Chemotaxis assays

Chemotaxis assays was performed as described previously [55] with minor changes. Briefly, L1 worms of strain CL2355 and their no-Aβ control strain CL2122 were incubated at 16°C for 36 hours on NGM plates containing 100 μg/mL ampicillin, and 1 mM IPTG seeded with *E. coli* containing either empty vector or vector that expresses double stranded RNA corresponding to the *dld-1* gene. The temperature was then up-shifted to 25°C for a further 36 hours. L4 stage worms were collected and washed with M9 buffer. After washing, worms were placed on the centre of the assay plate (with or without *dld-1* RNAi expressing *E. coli* lawn). Attractant containing 1 μl of odorant (0.1% benzaldehyde in 100% ethanol) was added in a spot on one edge of the plate with 1 μl of 100% ethanol as a control in a spot on the opposite side of the plate. 1 μl of 1M sodium azide was added to each of the two spots to immobilize the animals once they had migrated to one or the other destination. The chemotaxis index (CI) (number of worms at the attractant location - number of worms at the control location]/total number of worms on the plate) was calculated after 2 hours of incubation at 23°C.

### 2.10. Egg hatching assay

Wild type (N2), no Aβ control (CL2122) and the Aβ transgenic strain (CL2355) were synchronised and grown to maturity at 16°C (L4 stage, 4 days of age). 10 individuals were then transferred to fresh agar plates and temperature was shifted to 23°C. After 24 hours of incubation, adult worms were removed from plates. Unhatched eggs and larvae were counted every 24 hours for the next three days.

### 2.11. Uncoupler treatment

L4 worms were exposed to 17.5uM of the mitochondrial uncoupler carbonyl cyanide 4- (trifluoromethoxy)phenylhydrazone (FCCP). This dose does not cause significant mortality of wild type nematodes [56]. Mortality was scored immediately after a 24-hour exposure to FCCP at 23°C.

### 2.12. Oxidative stress measurement

#### 2.12.1. sod-3 expression

The response to mitochondrial superoxide-mediated oxidative stress was measured using *sod-3*::GFP in strain CF1553. Synchronized worms were fed with *E. coli* containing either empty vector or vector that expresses double stranded RNA corresponding to the *dld-1* gene for 72 hours at 20°C. Quantification of sod-3 levels was obtained using Zeiss fluorescence microscope. To get best signals, non-worm background fluorescence was subtracted from worm fluorescence.

#### 2.12.2 RO/NS measurement

Reactive oxygen/ nitrogen species (RO/NS) levels were measured using 2′,7′-dichlorofluorescein diacetate (DCF-DA) as described previously with modifications [57]. Briefly, worms were synchronized and placed on NGM plates with seeded with *E.coli* containing either empty vector or vector that expresses double stranded RNA corresponding to the *dld-1* gene. after 36 hours at 16°C followed by a temperature upshift for further 36 hours at 23°C, worms were washed with PBS three times and snap frozen in 250μl cell lysis solution. To prepare extracts, worms were sonicated followed by centrifugation at 14000rpm for 30 minutes. The supernatant was collected and further used for protein quantification using nanodrop. Supernatant containing 25μg of protein was pre-incubated with 250μM DCF-DA in 100μl of 1x PBS at 37°C for 1 hour. Fluorescence intensity (excitation wavelength 485nm and emission wavelength 535 nm) was measured using SpectraMax M3 fluoremeter (Molecular Devices, Sunnyvale, USA). To normalize fluorescent intensity, background fluorescence of 250μM DCF-DA was subtracted from each sample.

#### 2.12.3 H_2_O_2_ spectrophotometric measurement

Hydrogen peroxide (H202) levels were measured spectrophotometrically using toluidine blue as described previously by Sunil et al with minor modifications [58]. Worms extracts were prepared and quantified as described above in DCF-DA assay protocol. For each 25μg of protein we added 20μl 2% potassium iodide, 20μl 2M HCl, 10μl 0.01% toluidine blue and 40μl 2M sodium acetate. The contents were mixed and absorbance was measured at 628nm against known concentrations.

### 2.13. Quantitative RT-PCR

Wild type or Aβ expressing worms strain CL4176 were fed with empty vector or dld-1 RNAi after synchronization. After 36-hour initial feeding at 16°C, temperature was raised to 23°C for 48 hours and worms were collected for RNA extraction. Total RNA was extracted using the acid-phenol (Trizol) method and converted to single stranded cDNA using Invitrogen SuperScript cDNA synthesis kit following prescribed protocol. Gene specific primers were designed using NCBI Primer-BLAST as follows: *Aβ* forward primer CCGACATGACTCAGGATATGAAGT, *Aβ* reverse primer CACCATGAGTCCAATGATTGCA; *dld-1* forward primer GATGCCGATCTCGTCGTTAT, *dld-1* reverse primer TGTGCAGTCGATTCCTCTTG; *act-1* forward primer CGCTCTTGCCCCATCGTAAG, *act-1* reverse primer CTGTTGGAAGGTGGAGAGGG; *gpd-2* forward primer TTCTCGTGGTTGACTCCGAC, and *gpd-2* reverse primer AGGGAGGAGCCAAGAAGGTAAC. *Aβ* or *dld-1* mRNA levels in worms were quantified using Rotor Gene Q (QIAGEN) thermocycler. The PCR conditions were 95°C for 30 s followed 35 cycles of 95°C for 20 s (melting), 55°C for 30 s (annealing), and 72°C for 40 s (extension). For qPCR quantification, SYBR^®^ Green JumpStart™ ReadyMix™ (Sigma) was used. The relative gene expressions were monitored using the *gpd-2* or *act-1* genes by the 2^-ΔΔCt^ method.

### 2.14. Western Blotting of Aβ

Aβ was identified in *C. elegans* strains by immunoblotting after separation on a 16% Tris-Tricine gel. A standard western blotting protocol was used except that SDS was omitted from the transfer buffer. Briefly, synchronized L4 worms were incubated at 23°C for 48h and were then washed with distilled water and quickly frozen in liquid nitrogen. Flash frozen worms were either stored at -80°C or sonicated twice in ice cold cell lysis buffer (50 mM HEPES, pH 7.5, 6 mM MgCl_2_, 1mM EDTA, 75 mM sucrose, 25 mM benzamide, 1 mM DTT and 1% Triton X-100 with proteinase and phosphatase inhibitors 1:100 ratio). After sonication, the lysate was centrifuged at 10000 rpm to remove insoluble debris and total protein in the supernatant was measured using Pierce Coomassie (Bradford) protein assay kit (Thermo Scientific) on a NanoDdrop spectrophotometer. From each sample, 80-100 μg of total protein was precipitated with acetone and dissolved in Novex^®^ Tricine SDS sample buffer (LC1676, Invitrogen) by heating to 99°C for 5 minutes. Samples were subjected to gel electrophoresis at 100V for 2.5 hrs in separate cathode (100mM Tris, 100mM Tricine, 0.1% SDS, pH 8.3) and anode (0.2M Tris, pH 8.8) running buffers. Proteins were transferred onto nitrocellulose membranes by electroblotting in transfer buffer (35 mM glycine, 48 mM Tris (pH = 8.8) and 20% methanol) for 70 min at 100V and stained with Ponceau S (0.1% Ponceau S in 1% acetic acid) for 5 minutes following de-staining with 10% acetic acid (5 minutes) and washing under water 3 times or until smell of acetic acid was completely removed.

For Aβ, the membranes were blocked overnight in 5% skim milk at 4°C to prevent nonspecific binding of antibodies. The primary antibody staining was done using the Aβ monoclonal antibody 6E10 (Covance) at 1:1000 dilution in TBS (50mM Tris, 150mM NaCl, pH 7.6) containing 1% skim milk for 3-4 hours at room temperature following three washes with TBS-T five minutes each.

For DLD-1 detection, anti-lipoamide dehydrogenase antibody (ab133551) was used. Same procedure was repeated for DLD-1 detection except 5%BSA in 1X TBST was used for anti-body detection. Anti-mouse IgG alkaline phosphatase antibody produced in goat (A3562, Sigma), and anti-rabbit IgG alkaline phosphatase antibody produced in goat (A3687, Sigma were used as secondary antibody at 1:10000 dilution in TBS containing 1% skim milk or in 1% BSA in 1X TBST. Secondary antibody staining was done for 1 hour at room temperature. After washing the membrane with TBST, the proteins were detected using BCIP/ NBT substrate system (Sigma) or BCIP/ NBT kit (002209) from Lifetechnologies dissolved in 1M Tris (pH 9.0).

### 2.15. Statistical analysis

Differences due to treatments, strains and RNAi gene suppression were analyzed for statistical significance using GraphPad prism 6.0d. A p value less than 0.05 was considered statistically significant.

## 3. Results

The effect of metabolic rate on Alzheimer’s disease is an unresolved issue. While a decline in respiration rate is associated with both age of onset and the severity of AD, there are possible alternative explanations. It may either trigger the age-related increase in AD, or it may be a response that protects against the progression of AD. We use gene suppression of *dld-1* to directly test the effect of metabolic suppression in several different *C. elegans* models of Aβ pathology. Specifically, we tested the effect of *dld-1* gene suppression on nematodes that express Aβ either constitutively or with temperature induction in muscle cells or constitutively throughout the nervous system. The general experimental paradigm is to expose the nematodes to conditions known to result in Aβ toxicity and to determine whether genetic suppression of *dld* activity influences that toxicity. In our study *dld-1* RNAi effectively suppress the *dld-1* mRNA and subsequent protein expression (Supplementary Fig S1).

### 3.1. dld-1 suppression alleviates Aβ pathology in transgenic C. elegans

Transgenic expression and deposition of Aβ in body wall muscle cells of *C. elegans* causes severe, age-progressive paralysis. (Fig 1A). A temperature shift to 25°C was used to induce high level expression of Aβ. Fewer than 10% of the nematodes of the CL802 control strain that lacks the human Aβ transgene were paralyzed, i.e. unresponsive to prodding by 38 hours. In contrast, 100% of the worms of the CL4176 strain that does express human Aβ were unresponsive (Fig 1A). Suppression of *dld-1* in CL4176 reduced the frequency of paralysis due to Aβ expression to only ~30%, whereas suppression of the *dld-1* gene did not alter the robust activity of the control strain, CL802. When we extended the time of the assay (Fig 1B), we found that CL4176 worms in which *dld-1* gene expression had been suppressed did not become completely paralyzed until 144±24 hours. We repeated the test on CL2006 worms in which Aβ is expressed constitutively and found that suppression of the *dld-1* gene also delayed paralysis in these worms (Supplementary Fig S2). Thus, *dld-1* gene suppression prevents, to a large degree, the pathology associated with Aβ that causes paralysis.

**Fig 1:**
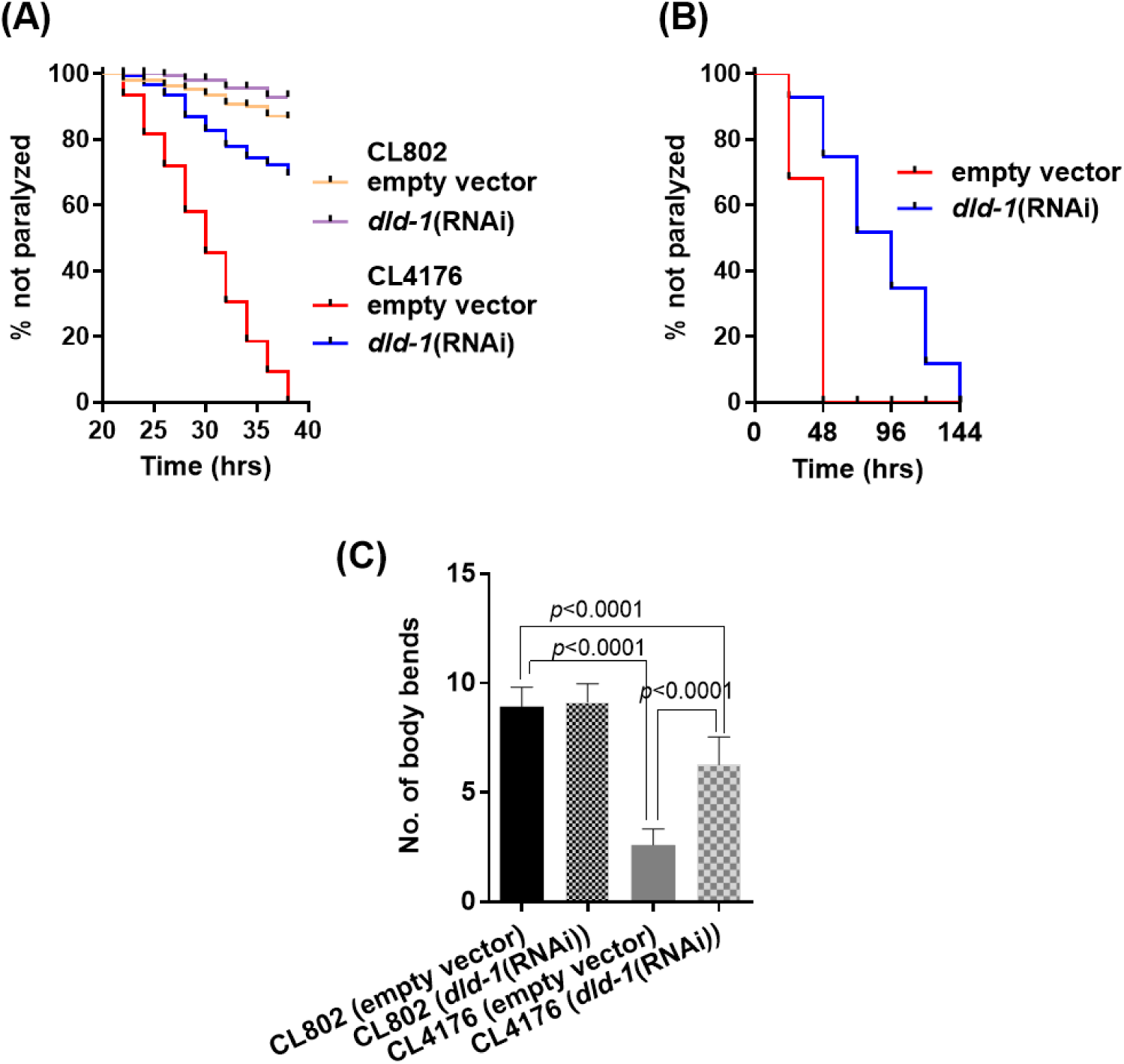
Suppression of the *dld-1* gene alleviates paralysis due to human Aβ expressed in transgenic *C. elegans.* (A) Time-dependent paralysis of the Aβ-expressing strain CL4176, with and without *dld-1* RNAi. Paralysis of synchronized L1 worms was measured on NGM plates seeded with *E. coli* strain HT115 containing either empty vector or a *dld-1* RNAi construct. After 36 hours at 16°C, the temperature was up-shifted to 25°C. Paralyzed worms were counted 24 hours after the temperature shift and thereafter, every two hours. (B) Extended time-dependent paralysis analysis. As *dld-1* RNAi significantly delayed paralysis, we repeated the analysis in a, but monitored the worms every 24 hours until the last worm become paralyzed. Kaplan Meyer survival curves were compared using a Logrank test. (C) CL4176 worms were synchronized and placed on plates either seeded with *E. coli* strain HT115 containing either empty vector or a *dld-1* RNAi construct. After 36-hour incubation at 16°C, the temperature was increased to 23°C for 36 hours. Worms were collected, washed and transferred to fresh plates. Worms were touched at the head region with a platinum wire and total number of body bends were counted under the microscope at 20°C. Results represented data from three independent trials (n=40-60 worms/ trial). Bars = mean ± SD.

Due to critical position in energy production cycle, *dld-1* suppression could result in reduction in glycolysis and downstream energy metabolism (TCA cycle and oxidative phosphorylation). We noted that addition of 5mM 2-deoxy-d-glucose (an analogue of glucose that does not enter TCA cycle thus reduce oxidative phosphorylation) to media also reduce *dld-1* mRNA and DLD protein expression and protect against Aβ-mediated paralysis in mutated worms (Supplementary Fig S1 and S3).

A second movement assay was performed that involved tapping the worms with a platinum wire and counting the number of body bends for 30 seconds. The worms were prepared as for the preceding assay except that the assay was carried out at room temperature (20°C) immediately after temperature induction of Aβ expression at 23°C. We found that as with the immobility assay, expression of human Aβ in the CL4176 strain resulted in a decrease in the rate of movement. Suppression of the *dld-1* gene by RNAi significantly improved mobility of CL4176 worms expressing human Aβ, resulting in 6.2±1.3 rather than 2.6±0.7 body bends (p<0.0001) (Fig 1C). The control CL802 worms that did not contain the Aβ transgene were unaffected by *dld-1* suppression (9.1±0.8 rather than 8.9±0.9 body bends) (p=0.529).

Expression of Aβ in muscle cells inhibits acetylcholine (ACh) neurotransmission, which is related to the observation that ACh agonists are commonly used to delay the symptoms of Alzheimer’s disease [59]. The inhibition of cholinergic neurotransmission by Aβ can be conveniently assayed by the protection it provides against a normally toxic dose of cholinergic agonist. Thus, restoration of normal sensitivity to the agonist is an indication of a decrease in the neurotoxic effects of Aβ.

To check whether *dld-1* inhibition restores normal ACh neurotransmission in CL2006 worms that constitutively express Aβ in muscle, we monitored paralysis in response to the cholinergic agonists, aldicarb (a potent acetylcholinesterase inhibitor) and levamisole (a cholinergic receptor agonist).

Resistance of the CL2006 strain to ACh agonists is due to production and deposition of both Aβ oligomers and fibrils [60]. Exposure to aldicarb (Fig 2A) results in paralysis within 180 min, which, as expected, occurs more rapidly under *dld-1* gene suppression (within 120 min, p=0.0001). Similarly, paralysis in response to levamisole is decreased from 240 min to 150 min (Fig 2B) when the *dld-1* gene is suppressed by RNAi (p=0.0001). Unlike the response in strains in which Aβ is expressed, suppression of *dld-1* had no effect on the response to either aldicarb or levamisole in either wild type or *dld-1* mutant worms (Supplementary Fig S4). Both nontransgenic strains became paralyzed earlier than transgenic worms that express Aβ (~120 min for both aldicarb and levamisole). Our results indicate that *dld-1* suppression restores near normal ACh neurotransmission in Aβ expressing worms via a decrease in Aβ-toxicity.

**Fig 2:**
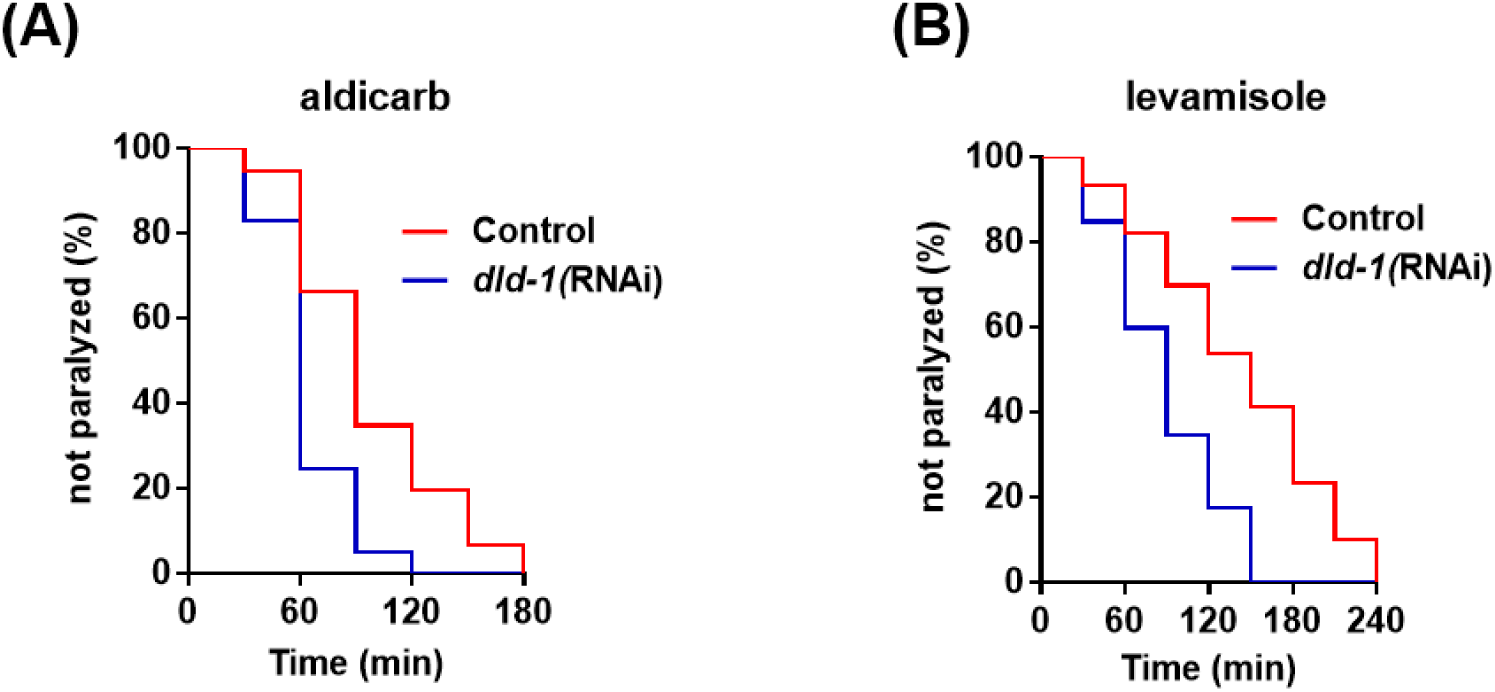
Acetylcholine neurotransmission assay in worms that express Aβ in muscle. Paralysis assay show that *dld-1* gene suppression improves acetylcholine neurotransmission in constitutive Aβ expressing *C. elegans* strain CL2006. (A) Time-dependent paralysis of control and transgenic worms fed on aldicarb (1 mM) with and without *dld-1* RNAi. (B) Time-dependent paralysis of worms fed on levamisole (0.2 mM) with and without *dld-1* RNAi. Results represented data from three independent trials (n=40-60 worms/ trial). Assay curves were compared using Log-rank test. Results represent the average of three independent trials.

Additional assays have been developed to monitor toxicity of *Aβ expressed in neurons*, impaired chemotaxis, hypersensitivity toward serotonin (5-HT), and reduced fecundity and egg hatching [43, 61]. Neural expression of Aβ in strain CL2355 significantly impaired chemotaxis toward benzaldehyde (chemotaxis index=0.05±0.01) relative to the non-Aβ control strain CL2122 (CI=0.20±0.02, p=0.002) (Fig 3A). Whereas suppression of the *dld-1* gene did not affect chemotaxis of the control strain (CI=0.22±0.02, p=0.4), it significantly improved chemotaxis of strain CL2355 (CI=0.14±0.01, p=0.002). Suppression of the *dld-1* gene in Aβ expressing worms was unable to fully restore chemotaxis to control levels (p=0.02).

Serotonin is an important biogenic amine neurotransmitter that mediates locomotion, egg laying and feeding behaviour in *C. elegans.* Exogenously applied serotonin causes paralysis in worms, which is exacerbated by expression of human Aβ [43]. In our study, 64±4% of worms of the control strain CL2122 were active, but this was reduced to 27±3% in the CL2355 strain that expresses Aβ throughout the nervous system (p=0.002). Suppression of the *dld-1* gene did not affect the activity of the control strain CL2122, (57±7%, p=0.3), but could partially alleviate serotonin induced paralysis in CL2355, increasing the percentage of worms that were active to (49±6%, p=0.004) but not to the level of the no-Aβ control strain (p=0.03). (Fig 3B).

**Fig 3:**
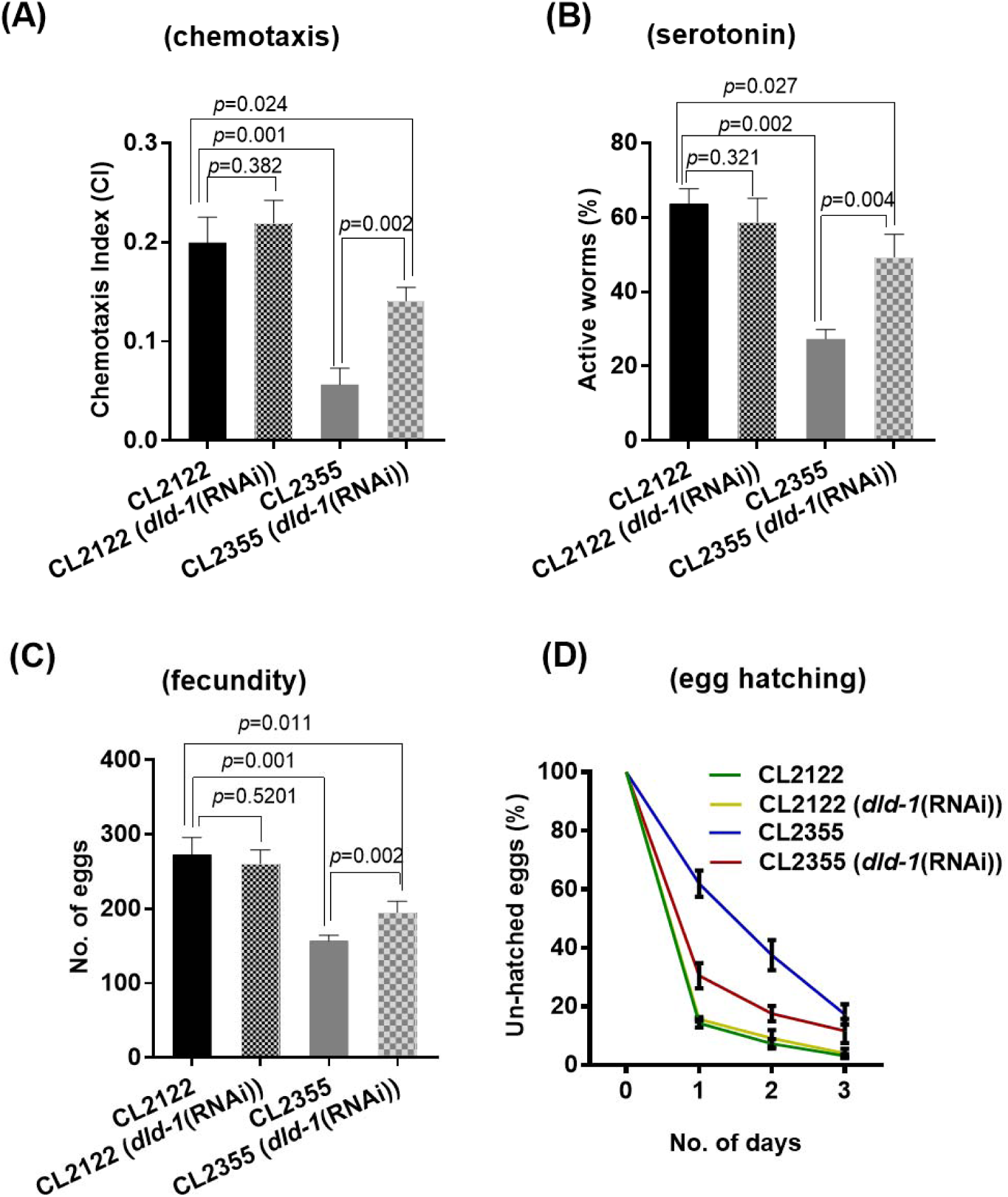
Effect of *dld-1* suppression on impaired behaviour in *C. elegans* that express Aβ in neurons. Chemotaxis, 5-HT serotonin sensitivity and, egg laying and hatching were compared between a no-Aβ control (CL2122) and that express Aβ in neurons (CL2355). Synchronized worms were fed with *E. coli* containing either empty vector or a vector that expresses *dld-1* dsRNA. (A) Analysis of chemotaxis behaviour in worms that express Aβ in neurons. Synchronized worms were placed at 16°C for 36 hours, and then shifted to 25°C. L4 worms were collected and assayed for chemotaxis towards benzaldehyde at room temperature (n = 40-50 worms in each well/ trial). (B) Evaluation of serotonin sensitivity in worms that express Aβ in neurons. Synchronized L4 stage worms were assessed for serotonin hypersensitivity at room temperature by placing them in a 96 well plate containing 250 μl of 1 mM serotonin (n = 25-30 worms / trial) and counted for paralysis after 5 minutes. Three independent trials were run for each experiment. (C) Fecundity of worms that express Aβ in neurons with or without *dld-1* RNAi. (D) Time course of egg hatching percentage in worms that express Aβ in neurons. For Fig 3C and 3D, worms were synchronized and placed at NGM plates with or without *dld-1* RNAi at 16°C until L4 stage appeared. Ten L4 stage worms were picked to fresh plates at 23°C to induce transgene expression. After 24 hours at 23°C, adults were removed. Plates were shifted to 20°C for remaining assay and eggs and larvae were counted each day for 3 days. Total number of eggs were estimated by adding the total number of un-hatched eggs and larvae present. Three independent trials were run for each experiment. Bars = mean ± SD.

Serotonin and ACh neurotransmission control egg laying [62], an activity that is inhibited by neuronal expression of Aβ. Based on our findings above, we reasoned that suppression of the *dld-1* gene would reverse the negative effect of Aβ expression on fecundity. Aβ expression significantly reduced egg laying in CL2355 relative to the control strain, CL2122 (157±8 vs 273±23, p=0.001). While there was no significant effect of *dld-1* gene suppression by RNAi on the strain that did not express Aβ (273±23 vs 261±19, p=0.5), suppression of the *dld-1* gene caused a marked improvement in fecundity in CL2355 (157±8 vs 207±11, p=0.002). The improvement in fecundity did not reach that of the matched control (p=0.01) (Fig 3C).

Aβ expression also negatively affects egg hatching, with 61.8% of CL2355 eggs remaining un-hatched after 24 hours. In contrast, only 14.2% of CL2122 eggs remained unhatched. Inhibition of *dld-1* resulted in a significant decrease in unhatched eggs after 24 hours, 30.5% (Fig 3D). The same trend persisted over the next two days. There was no effect of *dld-1* suppression on egg hatching of the control strain CL2122.

### 3.2. dld-1 suppression reduces Aβ protein oligomerization without affecting Aβ peptide levels

A reduction in Aβ toxicity in our study could result from either a decrease in overall *Aβ* peptide levels or a decrease in the formation of toxic oligomers [63]. We did not observe any significant change in *Aβ* mRNA levels after *dld-1* suppression, indicating that gene expression was not affected (Fig 4A). We then assessed whether *dld-1* suppression affected either the total amount of *Aβ* peptide produced or the degree of *Aβ* oligomerization. We found no change in the overall level of Aβ peptide due to *dld-1* suppression (Fig 4B and 4C). However, there was a significant decrease in the proportion of Aβ peptide in the form of ~19kDa oligomers and a corresponding increase in ˜4kDa monomers (0.31±0.13 vs 0.71±0.023, p=0.045) when the *dld-1* gene was suppressed in strain CL4176 (Fig 4B and 4D). In contrast, there was no significant change in oligomers of 12kDa, 16kDa or 23kDa.

**Fig 4:**
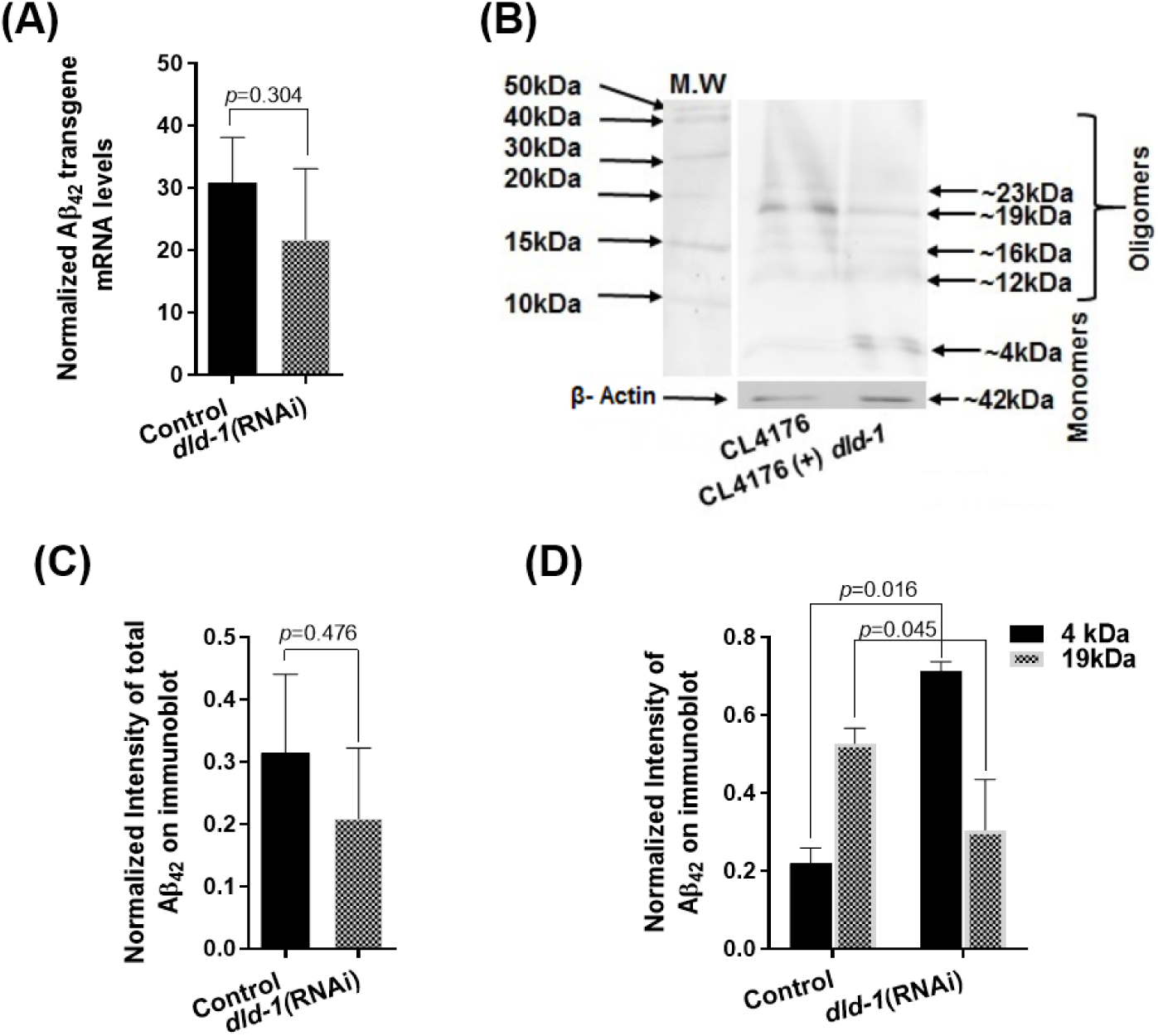
Effect of *dld-1* suppression on *Aβ* transgene and protein expression. (A) Quantitative RT-PCR of Aβ mRNA levels Synchronized worms were fed with *E. coli* containing either empty vector or a vector that expresses *dld-1* dsRNA. Worms were incubated for 36 hrs at 16°C and then temperature was shifted to 23°C for further 36 hrs before worms were collected for RNA or protein extraction. Levels of *Aβ* mRNA are normalized to glyceraldehyde-3-phosphate dehydrogenase (*gpd-2*) transcript levels, with experiments replicated three times. (B) A western blot of total soluble protein run on a 16% Tris-tricine gel shows multimers of Aβ *in C. elegans* strains expressing human Aβ. Arrows indicate the presence of *Aβ* monomers at 4kDa, and oligomers at 12kDa, 16kDa, 19kDa and 23kDa. (+) *dld-1* indicates treatments in which the *dld-1* gene has been suppressed by RNAi. The control using an anti-actin antibody (ab14128) is shown below to indicate the relative amount of protein loaded onto each lane. (C) Densitometry of overall protein bands appeared on western blot of each column to estimate differences in Aβ protein expression after *dld-1* suppression. (D) Quantification of the intensity of 4kDa monomer and 19kDa oligomer bands with and without *dld-1* suppression from three independent trials. Quantification was carried out using GelQuantNET software. Bars= mean±SD.

**Fig 5:**
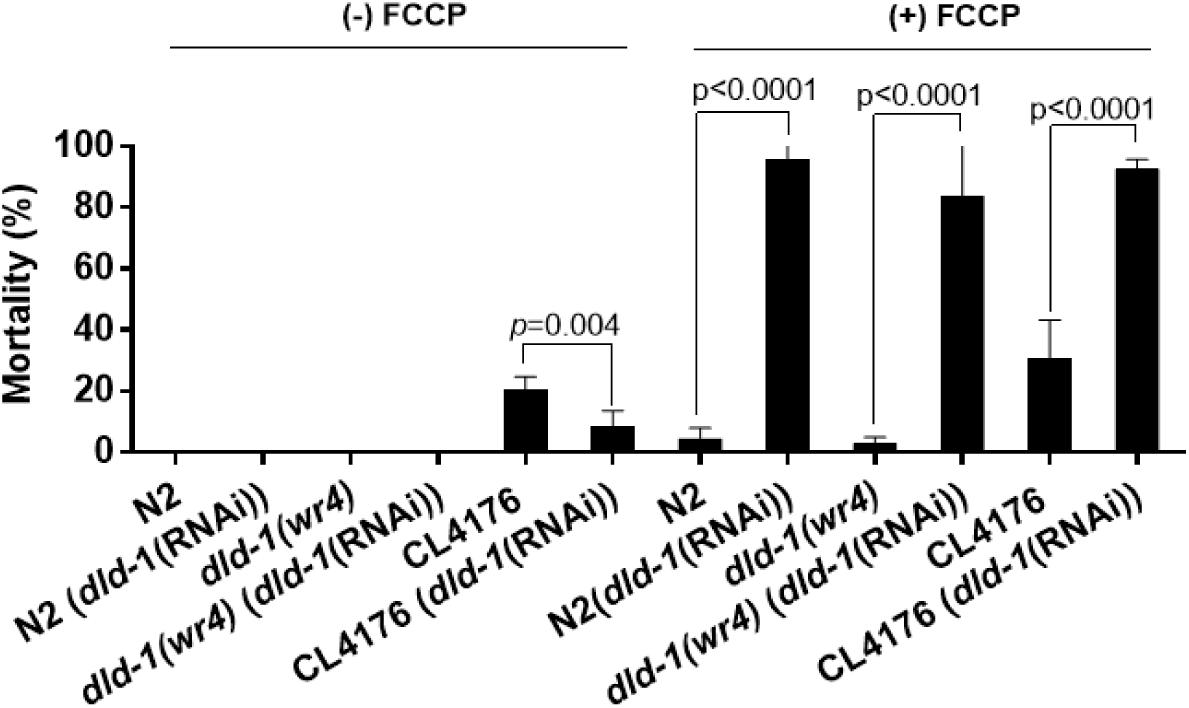
Mortality assay of *C. elegans* treated with the mitochondrial uncoupler FCCP. When the *dld-1* gene was suppressed in the presence of uncoupler, the compounded disruption of energy metabolism resulted in high level mortality in each of the three strains regardless of the presence of the *Aβ* peptide (n= 50-100 for each experiment. Three independent trials were run. Bars= mean ± SD.

### 3.3. How does dld-1 gene suppression protect against the toxicity of Aβ?

DLD-1 is a core enzyme of oxidative respiration and the *dld-1*(*wr4*) mutation is known to inhibit energy metabolism (Zuryn et al., 2008). FCCP is also a disruptor of mitochondrial energy metabolism, but it acts in quite a different manner. Both likely disrupt the generation of ATP, but by opposite mechanisms. DLD-1 disruption slows the flow of metabolites through the TCA cycle and the electron transport chain, whereas FCCP accelerates the same flows, but in a futile manner. Exposure to FCCP decreases Aβ production[64, 65], which implies that ATP depletion is more important to the protection than is the mechanism that causes the decrease in ATP.

To determine which of these two options, metabolite flux or ATP generation is more likely responsible for the protection against Aβ-induced toxicity, we measured Aβ-mediated toxicity in combination with either *dld-1* gene suppression or FCCP exposure or both. In the absence of FCCP, both wild type and *dld-1* mutant strains exhibited 100% survival whether or not *dld-1* was suppressed. The Aβ transgenic worms exhibited moderate mortality that was significantly alleviated by RNAi suppression of *dld-1* (20.5±4.2 vs 8.2±5.3, p=0.004).

Exposure to FCCP had an effect very different to that of *dld-1* gene suppression. The dose of FCCP that was used caused negligible mortality on its own, but rather than protect against Aβ-mediated mortality, it produced an apparent, but not quite significant increase in mortality from 20.5%±4.2 to 30.8%±12.3 (p=0.107). This outcome does not allow us to attribute protection against Aβ to a decrease in ATP generation. Mortality due to Aβ expression combined with FCCP increased greatly when *dld-1* gene expression was suppressed by RNAi (30.8±12.3 vs 92.9±2.8, p<0.0001). Mortality was not restricted to the transgenic strain, however, but also was observed in wild-type N2 (4.2±3.5 vs 95.7±4.1, p<0.0001) and the *dld-1*(*wr4*) mutant (3.7±3.3 vs 83.2±16.8, p<0.0001). The most likely explanation is that the decrease in metabolite flux due to a decrease in DLD-1 containing metabolic complexes, together with futile pumping of protons across the inner mitochondrial membrane caused by FCCP, results in a crisis of energy metabolism that affects all three strains equivalently, and that this is largely independent of whether Aβ peptide is expressed.

Another possible mechanism whereby *dld-1* gene suppression could reduce the toxicity of Aβ is via a decrease in reactive oxygen/ nitrogen species (RO/ NS), as ROS can induce aggregation of Aβ [24, 66, 67]. DLD itself can generate significant amounts of ROS (superoxide), so suppression of DLD activity could lead to a decrease in superoxide production [68, 69]. The superoxide dismutase-3 enzyme (SOD-3) converts superoxide into O_2_ and H_2_O_2_. Because the *sod-3* gene is induced by its substrate, superoxide, we used a strain of *C. elegans* (CF1553) that expresses GFP under the control of the *sod-3* promoter to measure the effect of *dld-1* suppression on intracellular superoxide levels. Suppression of the *dld-1* gene resulted in a decrease in GFP signal (28120.3±7884.3 vs 16662.8±6145.7, p=0.0016) (Fig 6A).

**Fig 6:**
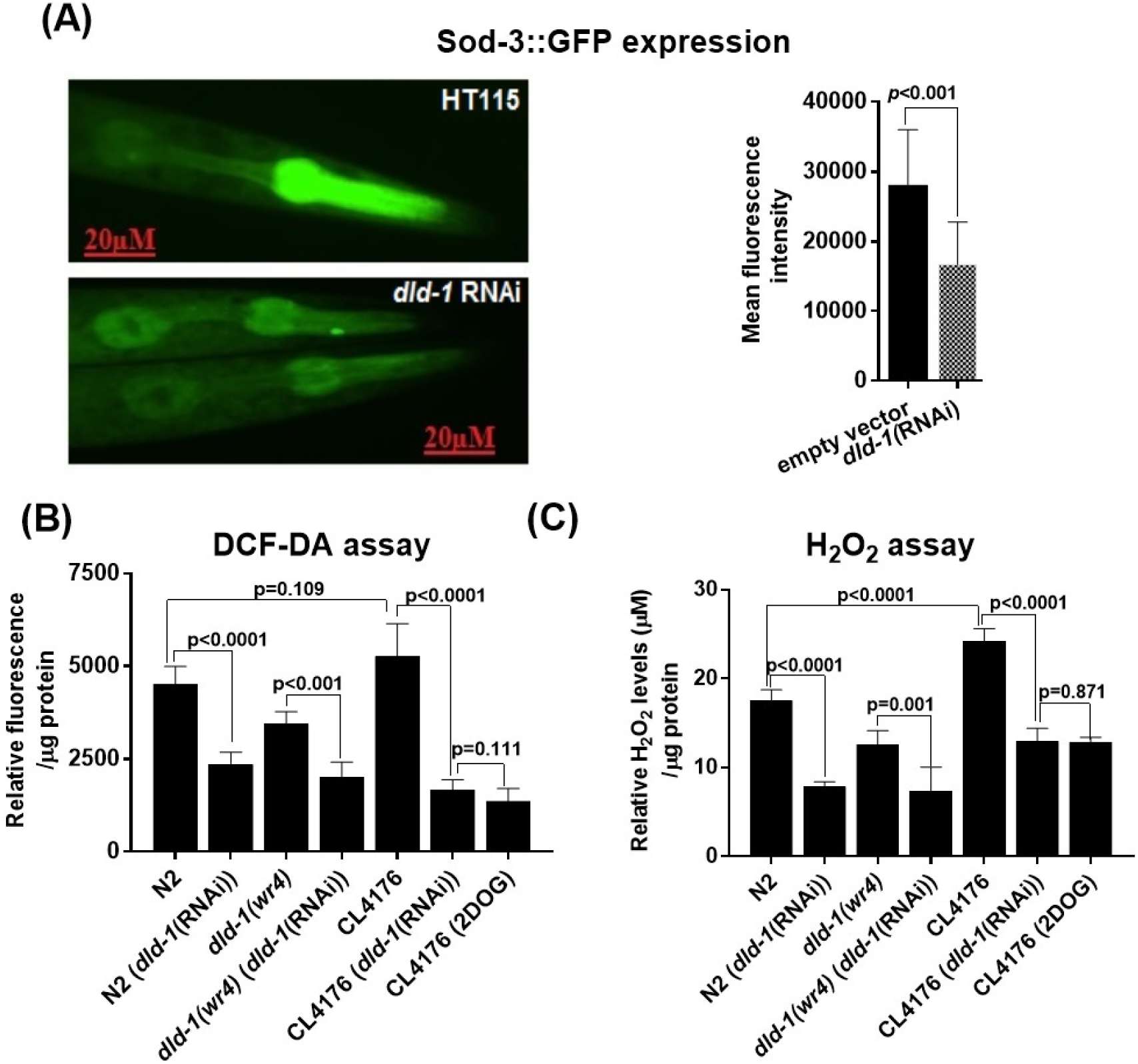
Suppression of the *dld-1* gene lowers the response to superoxide and lower RO/NS burden in *C. elegans.* (A) Synchronized L1 worms of strain CF1553 that expresses *sod-3*::gfp was fed *E. coli* containing either empty vector (strain HT115) or vector that expressed *dld-1* ds-RNA for 72 hours at 20°C. GFP fluorescence was quantified in at least ten worms from each group. Mean fluorescence intensity was measured using the formula; Integrated density - (area of selected worm × mean fluorescence of background readings). (B) Measurement of R)/NS levels in worms using DCF-DA. (C) Measurement of H_2_O_2_ levels in worms. Worms were synchronized and placed on NGM plates seeded with *E. coli* containing empty vector or expressing *dld-1* ds-RNA, and/ or containing 5mM 2DOG. Bars = mean ± SD.

**Fig 7:**
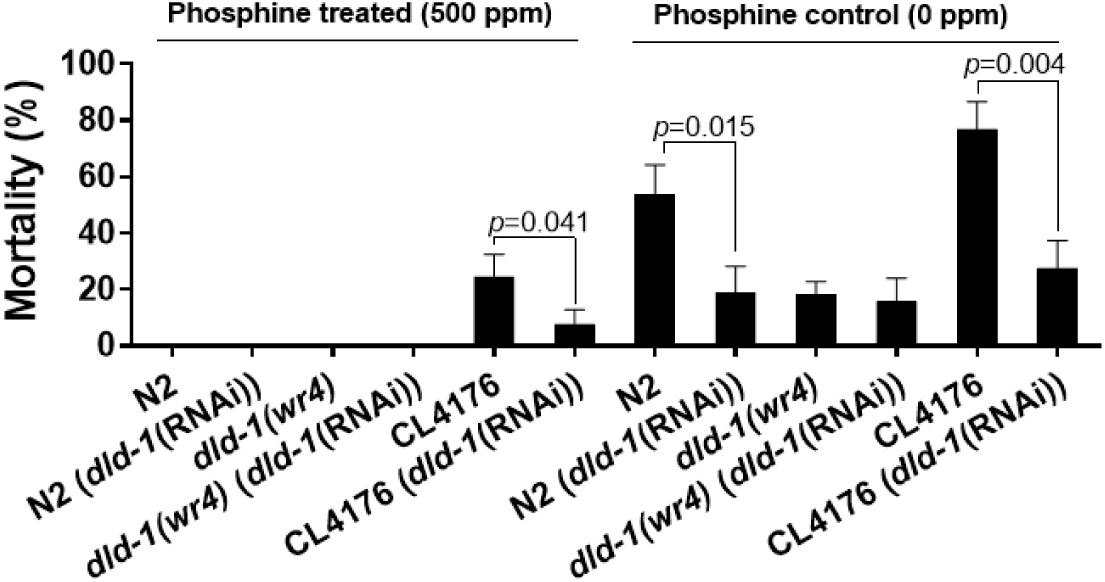
Suppression of the *dld-1* gene protects against the toxicity of phosphine independent of Aβ presence. *dld-1* gene mutation, or suppression by RNAi, reduced mortality caused by 500 ppm phosphine in both wild type (N2) and Aβ-expressing worms of strain CL4176. Three independent trials were run (n=50-80 for each trial). Bars = mean ± SD.

To further elaborate our observation, we measured the cellular RO/NS levels using DCFDA. Our results showed that *dld-1* mutant worms have lower RO/NS levels when compared to wild-type (3455±322vs4527±472, p=0.001). *dld-1* suppression significantly decreased the RO/NS levels in the worms regardless of phenotype. *dld-1* suppression not only decreased the RO/NS levels in wild type (4527±472 vs 2359±329, p<0.0001) and Aβ expressing strain (5266±883vs1675±262, p<0.0001) but also in *dld-1* mutant (3455±322vs2023±395, p<0.001). Interestingly, we observed no difference of RO/NS levels between wild type and Aβ expressing strain CL4176 (4527±472vs5266±833, p=0.109). Treatment with 2DOG also reduced the oxidative stress in Aβ expressing worm that was like the *dld-1* suppression (1675±333vs1371±335, p=0.111) (Fig 6B).

As described earlier DLD is a major source of superoxide production, this superoxide readily converted into hydrogen peroxide (H_2_O_2_). Quantification of H_2_O_2_ could be a good indicative of DLD activity and oxidative stress too. Although DCF-DA has been previously used for H_2_O_2_ measurement, recent data showed that DCF-DA does not react with H_2_O_2_ and does not form fluorescent product, and hence cannot be used for H_2_O_2_ quantification [70]. To overcome this limitation, we directly measure the H_2_O_2_ levels in the worm’s extracts spectrophotometrically. *dld-1* suppression significantly lower the H_2_O_2_ levels in wild type (17.5±1.2 vs 7.8±0.6, p<0.0001), *dld-1* (12.5±1.6 vs 7.3±2.6, p=0.001) mutants and Aβ expressing worms (24.1±1.4 vs 12.9±1.5, p<0.0001). We observed higher H_2_O_2_ levels in wild type when compared to *dld-1* mutant (17.5±1.2 vs 12.5±1.6, p<0.001). It is worth noting that H_2_O_2_ levels were significantly higher in Aβ expressing worms than wild-type (17.5±1.2 vs 24.1±01.4, p<0.0001). Treatment with 2DOG also decreased the H_2_O_2_ levels in Aβ expressing worms (24.1±1.4 vs 12.8±0.5, p<0.0001).

Our results indicate that suppression of *dld-1* gene expression causes a decrease in ROS, provide another plausible mechanism to explain the protective effect of *dld-1* gene suppression against Aβ-toxicity.

The *dld-1*(*wr4*) mutation that is used in the current study confers resistance against phosphine toxicity [71] a phenotype that can also be achieved by *dld-1* gene suppression[45]. Phosphine is a fumigant that induces ROS production and lipid peroxidation but causes decreased respiration rates, as well as a reduction in mitochondrial membrane potential and ATP levels [72]. Thus phosphine, like Aβ, impairs mitochondrial function causing phenotypes that are countered by *dld-1* inhibition. Due to these similarities, we investigated interactions between *dld-1*, phosphine and Aβ.

We found that exposing Aβ expressing transgenic worms to 500 ppm phosphine (the LC_50_ of wildtype *C. elegans*) for 24 hours following by 48 hours of recovery at room temperature increased the mortality of the wildtype N2 strain to the same degree as the Aβ expressing strain CL4176. Thus, mortality of N2 increased from 0% to 48.5±10.7% in response to 500ppm phosphine and mortality of CL4176 increased from 22.9±7.8% to 75.6±9.9%. The resistance phenotype of the *dld-1*(*wr4*) mutant was unaffected by RNAi-mediated suppression of the *dld-1* gene, indicating that the mutation and gene suppression confer resistance to phosphine by the same mechanism. The phosphine toxicity and Aβ toxicity is simply additive, regardless of whether or not the *dld-1* gene is suppressed. This indicates that while both phenotypes are affected by the DLD-1 enzyme, they are mediated independently without any interaction. Similar findings were observed when we treated the worms at a higher phosphine concentration of 2000ppm (Supplementary Fig S5).

## 4. Discussion

Imaging of the AD brain, consistently reveals a decrease in glucose metabolism in areas most strongly affected by disease pathology. While the effect of reduced glucose metabolism on AD progression has not been determined, the deposition of Aβ and reduction in glucose metabolism both begin decades before any clinical manifestation of the disease, indicating a possible causal relationship between the two [73].

To assess the role of energy metabolism in AD-associated Aβ proteotoxicity in worms, we used RNAi to suppress the activity of the *dld-1* gene, which is known to suppress aerobic respiration [74, 75]. The DLD enzyme is also a subunit of two mitochondrial enzyme complexes, pyruvate dehydrogenase (PDH) and alpha-ketoglutarate dehydrogenase (KGDH). These complexes contribute to the oxidative respiration of glucose and are implicated in AD as well [31, 76]. Our results show that suppression of the *dld-1* gene significantly alleviates the symptoms associated with Aβ expression in either muscles or neurons of *C. elegans*.

We find that suppression of the *dld-1* gene does not affect either the *Aβ* transgene mRNA levels or the levels of Aβ peptide. Suppression of *dld-1* does, however, significantly inhibit the oligomerization of Aβ. Accumulation of Aβ oligomers is thought to be a major culprit in AD progression [77, 78], whereas monomers actually help to maintain glucose homeostasis and are not toxic [79, 80]. Our findings suggest that *dld-1* suppression reduces Aβ oligomerization thus resulting in reduced paralysis, better movement rates, and improved behavioral phenotypes as observed previously [43, 60, 81, 82].

Recent studies have observed that diet rich in carbohydrates accelerates neurodegeneration by increasing Aβ oligomers [83, 84]. Suppression of the *dld-1* gene causes a decrease in glucose catabolism, which would appear to have the opposite effect of a carbohydrate-rich diet. In fact, dietary restriction has been shown to delay Aβ-mediated proteotoxicity and extend lifespan in *C. elegans* by activating the DAF-16/ FOXO, AAk-2/AMPK and SIR-2.1/sirtuin pathways [85-89]. Both mutation and RNAi-mediated suppression of the *dld-1* gene result in phosphine resistance and an extended lifespan (Kim and Sun, 2007; Cheng et al., 2003; Schlipalius et al., 2012), as well as inhibition of Aβ oligomerization and protection against Aβ-mediated toxicity as we have shown here.

Based on our understanding of the relationship between the *dld-1* gene and phosphine toxicity/resistance, we carried out several additional assays designed to compare the mechanisms of action of *dld-1* and Aβ. When we exposed worms to FCCP, we found that a dose that had no effect on either wildtype or *dld-1*(*wr4*) mutant worms and only a modest effect on Aβ expressing worms was highly toxic when the *dld-1* gene was subjected to RNAi-mediated suppression. The basis of the interaction between FCCP and *dld-1* gene suppression is unknown. FCCP does, however, deplete the mitochondrial proton gradient that is utilized for ATP synthesis, whereas the DLD enzyme generates NADH that delivers electrons to the electron transport chain that generates the proton gradient. It may be that simultaneous depletion of the proton gradient by FCCP as well as the source of electrons (NADH) by suppressing the *dld-1* gene, results in a cellular energetic catastrophe.

As protonophores, mitochondrial un-couplers also reduce the pH of the mitochondrial matrix. At low pH, the dehydrogenase activity of DLD is inhibited and the reverse activity (diaphorase) is induced [90, 91]. This would have the same effect on cellular energy metabolism as described in the previous paragraph and indeed, both mechanisms may contribute to the synergistic increase in mortality that is observed.

Aβ proteotoxicity and oxidative stress are positively correlated [67, 92, 93] and DLD inhibition is known to reduce ROS generation [68, 94, 95]. The DLD enzyme and mitochondrial electron transport chain (ETC) are both major sources of ROS generation [69, 96], so ROS production could either be direct (less ROS emanating from DLD) or indirect (less NADH feeding electrons to the ETC). We found reduced levels of the mitochondrial superoxide detoxifying enzyme SOD-3 after *dld-1* gene suppression, suggesting that suppression of *dld-1* does indeed decrease the burden of ROS in *C. elegans*.

We also exposed worms to the fumigant phosphine, a mitochondrial poison that causes oxidative stress and inhibits respiration, likely by targeting the DLD enzyme [45, 72]. When combined, Aβ expression and exposure to phosphine cause an additive increase in mortality. Suppression of the *dld-1* gene protects against both Aβ and phosphine individually and provides the same degree of protection against the two stressors in combination. One interpretation of these results is that the two stressors act through the same mechanism(s), with each stressor simply increasing the magnitude of the insult.

The similarities that we observe between the toxicity of Aβ and phosphine are worth noting. Both cause suppression of energy metabolism and yet are protected by suppression of the *dld-1* gene, a manipulation that likewise suppresses energy metabolism. The toxicity of both Aβ and phosphine is synergistically exacerbated by co-exposure to the mitochondrial uncoupler, FCCP. However, co-exposure to Aβ and phosphine does not result in an additive rather than synergistically increased toxicity, an outcome that is consistent with Aβ and phosphine acting by the same mechanism.

Our findings establish a neuroprotective effect resulting from suppression of the *dld-1* gene that encodes a core enzyme of mitochondrial energy metabolism in *C. elegans* strains that express human Aβ. Although it is not clear how *dld-1* suppression regulated the Aβ oligomerization in our study, one depiction could be the change in calcium influx. There are some observations that induced calcium influx promote Aβ oligomerization and vice versa [91, 97]. DLD complex could be involved in regulating calcium homeostasis as induced H_2_O_2_ levels were found to increase internal calcium stores [98]. We find no evidence that metabolic suppression through DLD is a risk factor for Aβ proteotoxicity. Our results do not distinguish between two possibilities; that neuroprotection is a direct effect of metabolic suppression or that it is an indirect effect resulting from decreased ROS generation. Regardless of the mechanism, our results are consistent with the hypothesis that a decrease in mitochondrial energy metabolism protects against Aβ pathogenicity, which in humans could delay clinical dementia resulting from AD.

## Supplementary information

**Fig S1:**
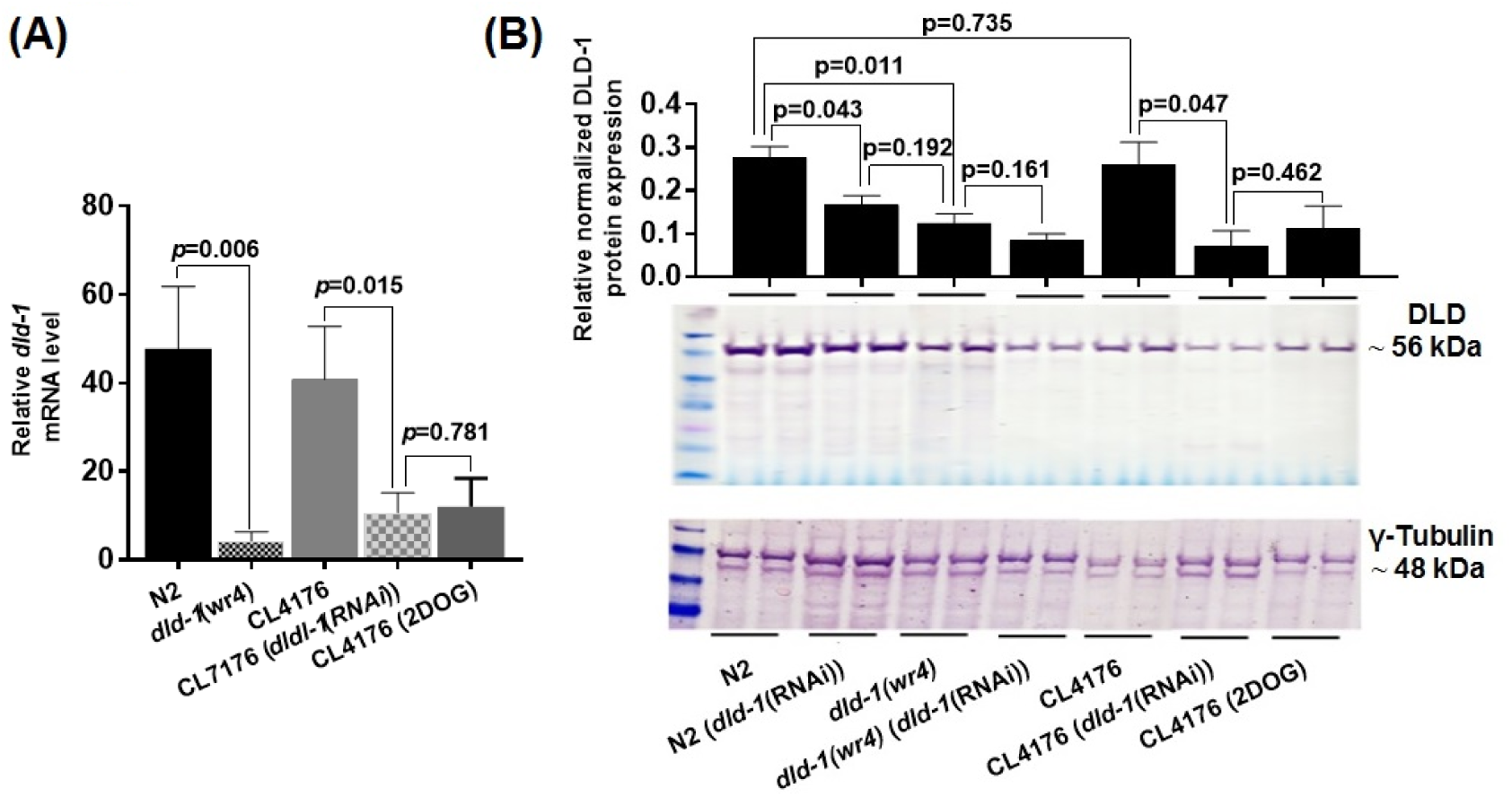
*dld-1* mutation, or suppression by RNAi causes a decrease in *dld-1* transcript and protein levels. Synchronized L1 stage worms of the Aβ expressing strain CL4176 were fed *E. coli* that expressed *dld-1* dsRNA for 36 hours at 16°C. The temperature was raised to 25°C for 36 hours to enhance Aβ expression. Temperature was also increased to 25°C in control worms (N2 and *dld-1*(*wr4*)) that do not express Aβ peptide. (A) Results of real-time quantitative PCR from three independent trials showing a significant decrease in *dld-1* mRNA levels in *dld-1* mutated and suppressed worms. (B) Western blot of protein extracted from cell lysate. Anti-lipoamide dehydrogenase antibody (ab133551) was used to detect DLD-1 protein, whereas anti-γ-tubulin antibody (ab50721) was used as reference control. Quantification of the DLD-1 bands from western blots using GelQuantNET software. Graphs and western blots represent the results from three independent experiments. Errors bars = mean±SD. Real-time quantitative PCR and western blotting were used to confirm the efficacy of RNAi-mediated *dld-1* gene suppression. We assessed the *dld-1* mRNA and protein levels in the Aβ expressing strain CL4176 before and after *dld-1* suppression, compared to the wild type strain N2 and the *dld-1* mutant *dld-1(wr4).* When the *dld-1* gene was suppressed by RNAi, both transcript and protein decreased to the levels in *dld-1(wr4*) mutant.

**Fig S2:**
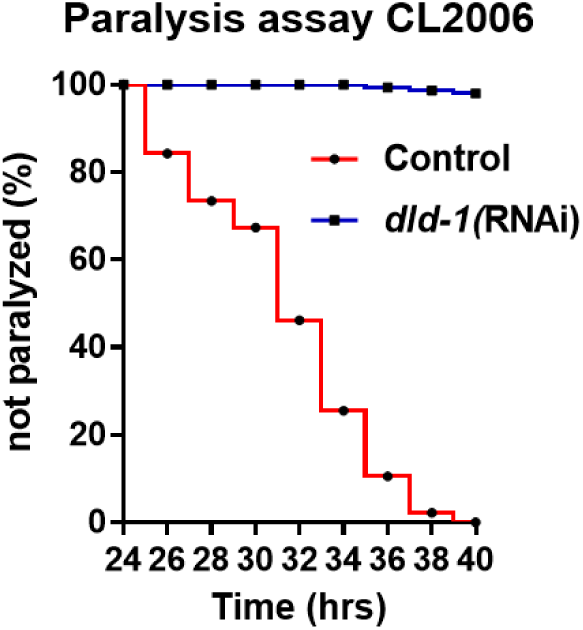
Effect of *dld-1* suppression on paralysis of the constitutively expressing Aβ strain CL2006. Synchronized worms were treated with *dld-1* RNAi for 36 hrs at 16°C. The temperature was shifted to 25°C for the next 24 hrs and scored for paralysis every 2 hrs. Paralysis curves were assessed using Log-rank survival test. Suppression of *dld-1* was found to delay the Aβ-associated paralysis in constitutively Aβ expressing worms just as occurred with the temperature inducible worms (Figure 1).

**Fig S3:**
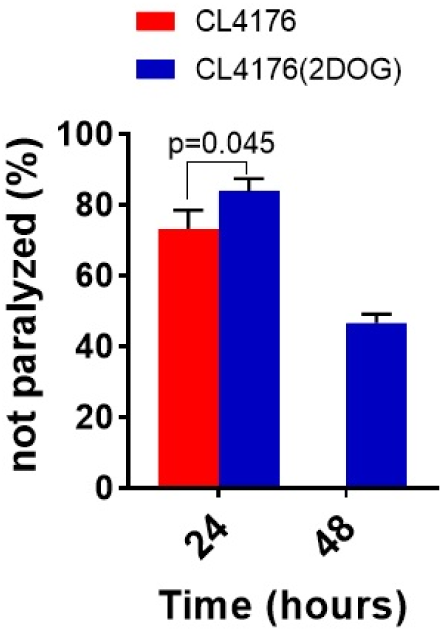
Effect of 5mM 2DOG on paralysis of the temperature inducible Aβ strain CL4176. Synchronized worms were placed on NGM plated with 5mM 2DOG for 36 hrs at 16°C. The temperature was shifted to 25°C for the next 24 hrs. Worms were scored for paralysis every 24 hrs. Addiction of 5mM 2DOG in media was found to delay the Aβ-associated paralysis in Aβ expressing worms just as occurred with the worms with suppressed *dld-1* expression (Figure 1). However, delayed paralysis was more profound in *dld-1* suppressed worms than treated with 2DOG.

**Fig S4:**
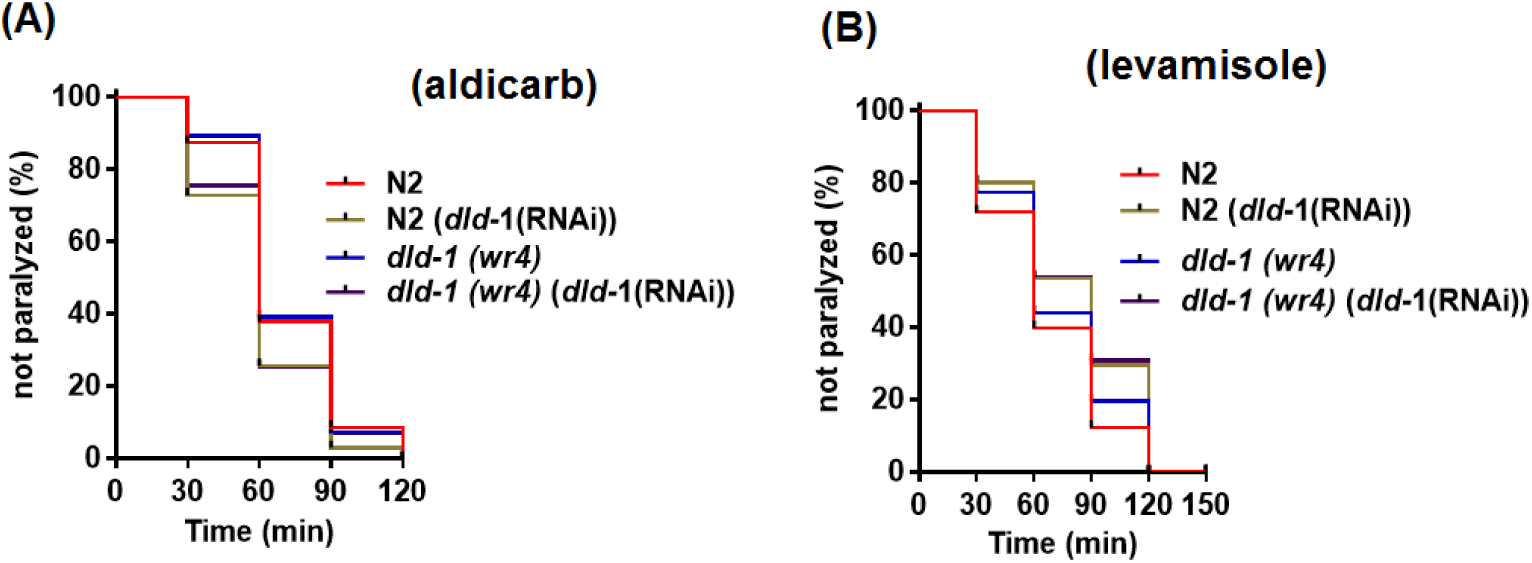
*dld-1* suppression does not change acetylcholine neurotransmission in wild type and *dld-1* mutated worms. *dld-1* was suppressed in Wild type N2 and the *dld-1* mutated worms using RNAi. After 72 hours of incubation at 20°C, worms were shifted to NGM plates containing either; (A) 1mM aldicarb or (B) 0.2mM levamisole. A log-rank test was applied to made comparison among treatments. The paralysis assay shows that *dld-1* gene suppression did not significantly affect acetylcholine neurotransmission in either wild type or *dld-1* mutant *C. elegans.* Thus, the improvement in acetylcholine neurotransmission observed in figure 2 was specific to the Aβ expressing strain, CL2006.

**Fig S5:**
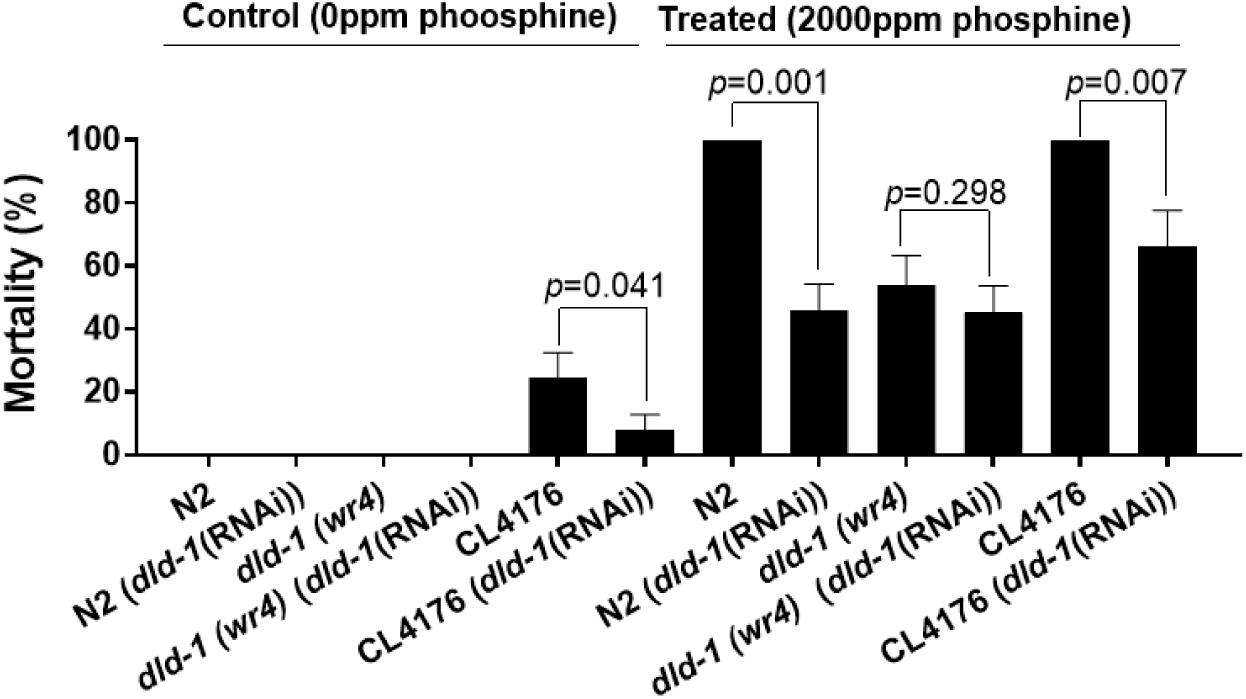
Suppression of *dld-1* gene protects against the toxicity of phosphine treatment at 2000ppm. Synchronized worms were either fed *E. coli* containing empty vector or expressing *dld-1* dsRNA. L4 stage worms were subjected to 2000ppm phosphine for 24 hours at 20°C. After a 48 hrs recovery time, worms were counted either dead or alive. Data are the result of three independent trials. Bars = mean± SD. Wild type N2 and *dld-1*(*wr4*) exhibited no mortality in the absence of phosphine. The Aβ expressing strain CL4176 exhibited mild paralysis that was reduced by *dld-1* suppression (22.9±5.9% vs 7.1±5.2%, p=0.041). Phosphine treatment at 2000ppm (LC_50_ of the *dld-1*(*wr4*) mutant) killed 100% N2 and CL4176 worms, but only 53.7±9.1% of the *dld-1* mutant worms. *dld-1* suppression decreased the mortality rate in N2 (54.5±8.2, p=0.001) and CL4176 (67.4±11.5, p=0.007), but there was no significant effect observed on *dld-1(wr4*) mortality (45.3±8.23, p=0.298).

